# Single-residue mutation in protein kinase C toggles between cancer and neurodegeneration

**DOI:** 10.1101/2023.03.16.532226

**Authors:** Alexander C. Jones, Alexandr P. Kornev, Jui-Hung Weng, Gerard Manning, Susan S. Taylor, Alexandra C. Newton

## Abstract

Conventional protein kinase C (PKC) isozymes tune the signaling output of cells, with loss-of-function somatic mutations associated with cancer and gain-of-function germline mutations identified in neurodegeneration. PKC with impaired autoinhibition is removed from the cell by quality-control mechanisms to prevent accumulation of aberrantly active enzyme. Here, we examine how a single residue in the C1A domain of PKCβ, arginine 42 (R42), permits quality-control degradation when mutated to histidine in cancer (R42H) and blocks downregulation when mutated to proline in the neurodegenerative disease spinocerebellar ataxia (R42P). Using FRET-based biosensors, we determined that mutation of R42 to any residue, including lysine, resulted in reduced autoinhibition as indicated by higher basal activity and faster agonist-induced plasma membrane translocation. R42 is predicted to form a stabilizing salt bridge with E655 in the C-tail and mutation of E655, but not neighboring E657, also reduced autoinhibition. Western blot analysis revealed that whereas R42H had reduced stability, the R42P mutant was stable and insensitive to activator-induced ubiquitination and downregulation, an effect previously observed by deletion of the entire C1A domain. Molecular dynamics (MD) simulations and analysis of stable regions of the domain using local spatial pattern (LSP) alignment suggested that P42 interacts with Q66 to impair mobility and conformation of one of the ligand-binding loops. Additional mutation of Q66 to the smaller asparagine (R42P/Q66N), to remove conformational constraints, restored degradation sensitivity to that of WT. Our results unveil how disease-associated mutations of the same residue in the C1A domain can toggle between gain- or loss-of-function of PKC.

## Introduction

Protein kinase C (PKC) is a family of multidomain Serine/Threonine protein kinases that propagate signals mediated by phospholipid hydrolysis (1–3). Activation of PKC isozymes leads to the regulation of diverse functions, including proliferation, cytoskeletal organization, and receptor internalization which position them as important players in the intracellular signaling landscape (4, 5). Dysregulation of PKC signaling output is evident in disease states, with impaired signaling generally associated with cancer (6) and enhanced signaling associated with neurodegenerative conditions such as Alzheimer’s disease (AD) (7, 8) and spinocerebellar ataxia type-14 (SCA14) (9).

PKC isozymes are designed to respond rapidly and reversibly to second messengers (2). They are matured by a series of ordered phosphorylations necessary to prime PKC into a stable, autoinhibited, but signaling-competent conformation ready to respond to second messengers. Without these phosphorylations, PKC is unable to adopt the stable autoinhibited conformation and is subjected to ubiquitin-mediated degradation (10, 11). For the subclass of conventional PKC isozymes (cPKC; α, βI, βII, γ), the first and rate-limiting phosphorylation is catalyzed by mTORC2 at the Tor Interaction Motif (TIM) and adjacent turn motif. These phosphorylations facilitate phosphorylation of the activation loop by phosphoinositide-dependent kinase-1 (PDK-1), in turn promoting autophosphorylation at the hydrophobic motif (12–17). This last event is necessary to lock cPKC into a stable and autoinhibited conformation by anchoring the C-tail to a conserved pocket on the N-lobe of the kinase domain (17–21). cPKC isozymes contain an N-terminal regulatory moiety that consists of a pseudosubstrate segment, diacylglycerol (DG)-sensing C1 domains, and a Ca^2+^-sensing C2 domain followed by the kinase domain and regulatory C-tail (2). Both C1 domains have been well-characterized for their ligand-binding properties (22–25), yet in the context of the fulllength protein the C1A domain is essential to maintaining PKC in an inactive state (26, 27) whereas the C1B is in an orientation favorable for binding DG and the functional analogues, phorbol esters. Reversible activation occurs via release of an autoinhibitory pseudosubstrate segment in response to allosteric activators (28), Ca^2+^ and DG, which are generated by the cleavage of membrane lipids by phospholipase C following GPCR activation (29). The Ca^2+^-bound C2 domain engages on the plasma membrane in an interaction involving phosphatidylinositol-4,5-bisphosphate (PIP2) (30), facilitating the binding of the C1B domain to its membrane-embedded ligand, DG, and other membrane lipids, including phosphatidylserine (23, 31). Engagement of these regulatory modules provides the energy to expel the pseudosubstrate from the active site to allow downstream signaling. In the open and active conformation, PKC is sensitive to dephosphorylation at the hydrophobic motif by the PH domain leucine-rich repeat protein phosphatase 1 (PHLPP1), ubiquitination, and proteasomal degradation (11, 32–34)—a process referred to as “downregulation.”

Alterations of cPKC isozymes in cancer are generally loss-of-function and occur by diverse mechanisms. These include reduced gene and protein expression (35), mutations that perturb processing phosphorylations, ligand binding, substrate binding, or catalytic activity (6), creation of fusion proteins that truncate the kinase domain or regulatory elements (32), or reduced protein levels due to a PHLPP1-mediaed qualitycontrol mechanism that degrades aberrantly active PKC (11). In contrast, gain-of-function mutants that evade or bypass quality-control degradation have been associated with neurodegenerative diseases. For example, one AD-associated mutation in PKCα (M489V) enhances the catalytic rate (k_cat_) of the enzyme without affecting the rate of activation and re-autoinhibition (k_on/off_) (8). This subtle mechanism allows the mutant protein to evade downregulation by maintaining autoinhibition in the absence of second messengers, but the signaling output is increased by 30% when activated. This small increase in activity is sufficient to cause cognitive decline in a mouse model, underscoring the importance of homeostasis in PKC signaling output (36). Gain-of-function mutations that evade downregulation are also associated with another neurodegenerative disease, spinocerebellar ataxia type-14 (SCA14). This autosomal dominant disease is caused by missense variants in *PRKCG*, the gene encoding PKCγ (37), and leads to cerebellar atrophy and loss of motor coordination and function (38, 39). Recent work has demonstrated that these mutants disrupt autoinhibition, leading to increased basal signaling output, yet are resistant to activator-induced degradation (9). Many of these mutated residues cluster to the C1 domains, including residues involved in ligand-binding or metal-coordination (22, 40), or are at predicted interfaces involving the C1 domains and catalytic domain of PKC (9, 41), suggesting that the integrity and/or interdomain contacts of the C1 domains are required for downregulation. Curiously, a highly conserved residue in the C1A domain (R42) is mutated both in cancer and in ataxia, but to different amino acids. R42 is mutated to histidine in cancer (42) and to proline in a family of patients diagnosed with SCA14 (PKCγ-R41P) (43). Understanding how mutation of a single residue can toggle between a cancer versus a degenerative phenotype would provide insight into how to target PKC in disease.

Here, we use biochemical, cellular, and *in silico* approaches to understand how disease-associated variants of a conserved arginine in the C1A domain of PKCβII, R42H (cancer), and in the equivalent position in PKCγ, R42P (SCA14), have opposing functional mechanisms. Whereas both mutations disrupt autoinhibition, the proline mutation uniquely disrupts the structure of the C1A domain, allowing the ataxia mutant to evade quality-control degradation. Our results provide a mechanism for disrupted signaling of these mutants and offer a structural explanation for how mutations at the same residue from two disease states can toggle between gain- and loss-of-function.

## Results

### R42 in C1A domain predicted to form ion pair with E655 in C-tail of cPKC isozymes

Due to the dynamic nature of PKC, a full-length, autoinhibited structure with all domains included and sufficient resolution has remained elusive; however, biochemical evidence has shown that the regulatory domains of PKC isozymes contribute to autoinhibition (44–47). Docking studies using individually crystalized domains (48, 49) and a partial structure of PKCβII (50) have been used to identify residues in the C1 and C2 domains that help maintain autoinhibition (51–53). Previously, we combined existing biochemical evidence and structural information to piece together a full-length model of the autoinhibited conformation of conventional and novel PKC isozymes (41). This model predicts an ion pair between R42 in the C1A domain and E655 in the C-tail, two conserved residues in cPKC isozymes, that appears to contribute to autoinhibition (Figure 1).

**Figure 1.**
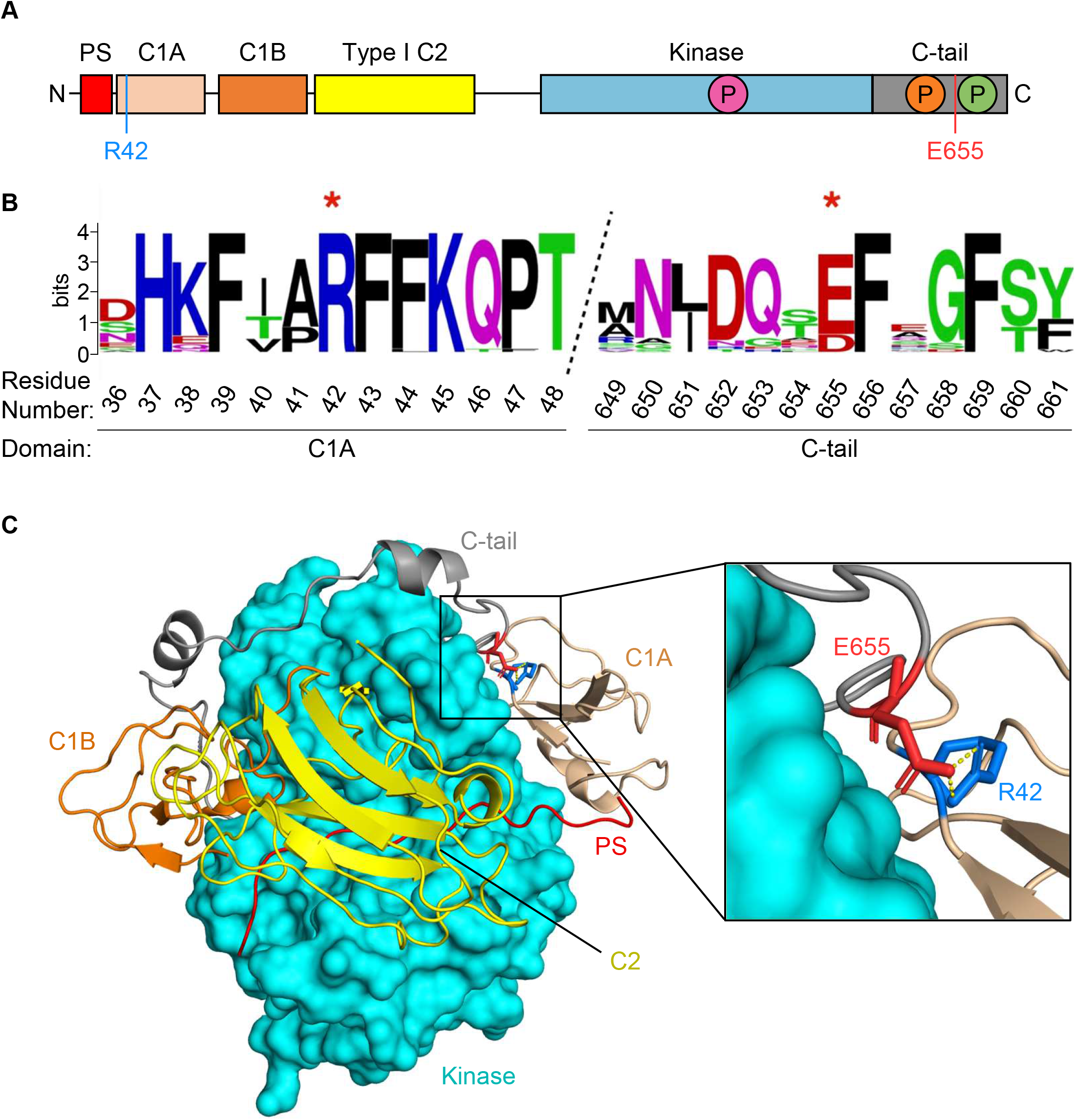
Conventional PKC isozymes contain a conserved salt bridge between the C1A domain and C-tail. (**A**) Domain structure of cPKC isozymes showing pseudosubstrate (PS, red) which reversibly occupies the active site, C1A domain (sand) and C1B domain (orange) which bind diacylglycerol, C2 domain (yellow) which acts as a Ca^2+^ sensor and binds phosphatidylserine and PIP2, kinase domain (cyan), and C-tail (grey). Three processing phosphorylations indicated in magenta, orange, and green for the activation loop, turn motif, and hydrophobic motif, respectively. Residues participating in the salt bridge are indicated in blue (R42) and red (E655). (**B**) Sequence logo showing conservation of R42 and E655 (starred) in bilaterian cPKC isozymes. Logo is drawn from alignment of representative sequences of vertebrate (one per class) and invertebrate (one per phylum) cPKC isozymes. R42 is absolutely conserved, and E655 is sometimes substituted by aspartate. (**C**) Structural model of PKCβII (41) in the autoinhibited conformation showing domains as colored in (A). Predicted ion pair shown between R42 and E655. Numbering corresponds to human PKCβII (UniProtKB: P05771-2).

Phylogenetic analysis revealed high co-conservation of R42 and E655 in higher order animals in cPKC isozymes (Figure 1B). R42 is conserved as an arginine in all bilaterian cPKC isozymes, with the exception of some very recent duplicate sequences. E655 is also absolutely conserved as glutamate or aspartate in all bilaterians, with the exception of a few recent duplicates and some alternative splice isoforms. This pattern suggests that the ion pair is functional in all bilaterians, while splice or duplication variants that have lost this regulation also occasionally emerge. cPKC isozymes are present in all holozoans, but in both choanoflagellates and sponges, the ion pair is not seen, and in cnidarians it is variably present and likely not key to function. These residues are mutated in multiple tumor types across the conventional family of PKC isozymes, including D652Y in PKCα, R42H and E655K mutations in PKCβII, and D669H/N in PKCγ (42). An R41P mutation in PKCγ (equivalent position to R42 in PKCβII) was also identified in a family of patients with SCA14 (43).

### Mutation of R42 or E655 disrupts autoinhibition of PKCβII

We previously demonstrated that mutation of E655 to lysine reduced autoinhibition of PKCβII as shown by more rapid plasma membrane translocation compared to WT in response to the C1 agonist phorbol 12,13-dibutyrate (PDBu) (52). To examine if R42 mutation also reduced autoinhibition (as our model predicts), we mutated this residue to alanine, glutamate, glycine, histidine, lysine, and proline and assessed the effect of each mutation on the rate of membrane translocation by monitoring FRET between YFP-tagged PKC and plasma membrane-targeted CFP (MyrPalm-CFP) (54, 55). All mutants translocated to the plasma membrane significantly faster than WT PKCβII, including PKC with a charge-conserved R42K mutation (Figure 2A-B). Our data are consistent with mutation of either R42 or E655 promoting a more open conformation of PKCβII.

**Figure 2.**
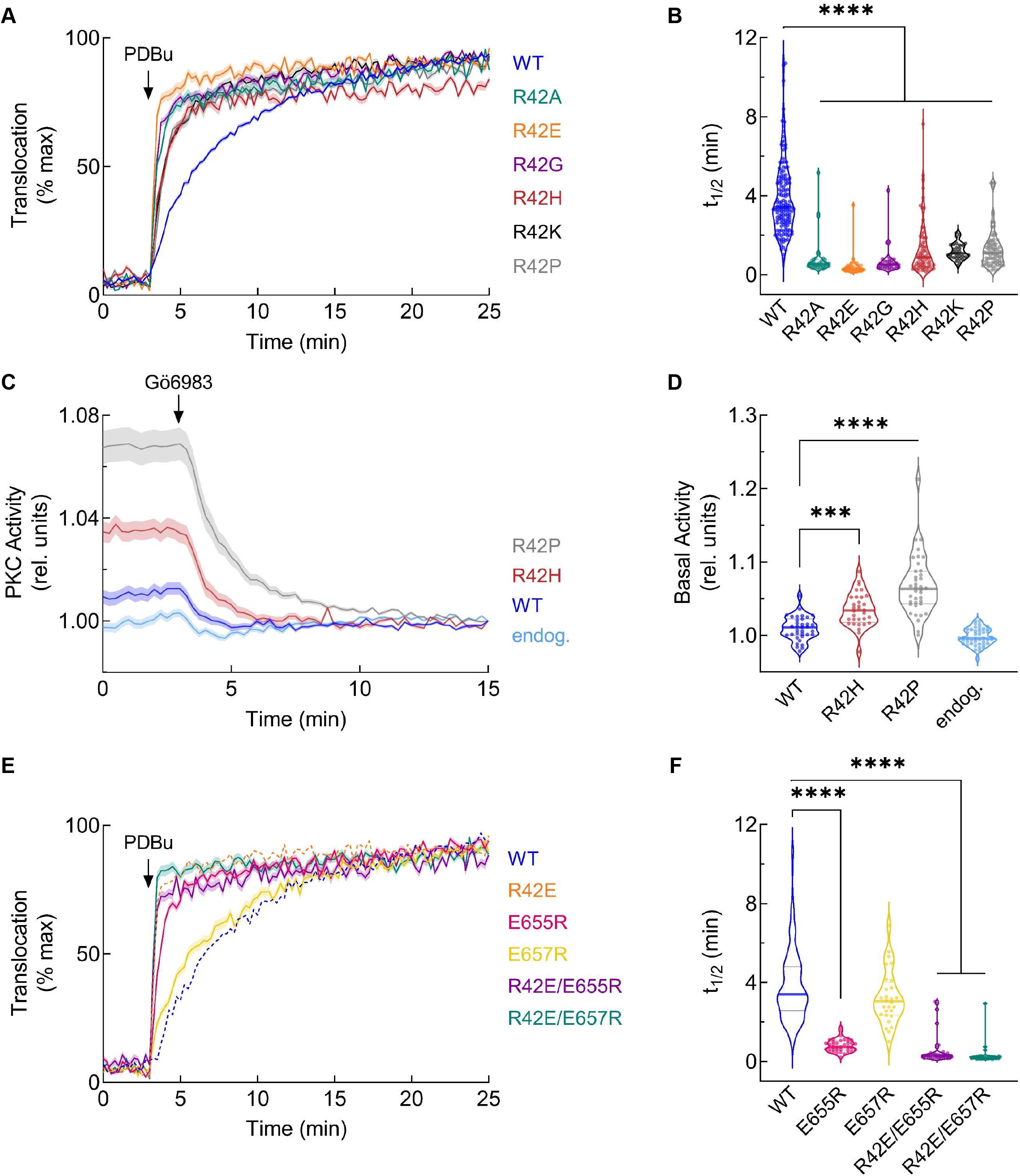
Translocation rate and activity of PKCβII-R42 mutants are increased in live cells. (**A**) COS7 cells were co-transfected with YFP-tagged PKCβII wild-type or mutants and MyrPalm-CFP (54). Translocation to plasma membrane was monitored by measuring FRET/CFP ratio changes after stimulation with 200 nM PDBu. Data for each cell were normalized to the max FRET ratio for that cell and represent at least three independent experiments and n ≥ 30 cells per condition. (**B**) Half-time of translocation was determined for each cell by fitting the data to a non-linear regression using a one-phase association equation. (**C**) COS7 cells were transfected with CKAR2 alone (endogenous, light blue) or co-transfected with indicated mCherry-tagged PKCβII construct. 1 μM of the PKC inhibitor Gö6983 was added after 3 min and PKC activity was monitored by measuring FRET/CFP ratio changes. Data were normalized to the assay end point and are from four independent experiments; n ≥ 32 cells per condition. (**D**) Basal activity from (C) was determined for each cell by plotting the initial FRET/CFP ratio after normalizing to the assay end point. (**E**) COS7 cells were co-transfected with YFP-tagged PKCβII and MyrPalm-CFP and plasma membrane translocation was measured as in (A). PKCβII-WT and R42E are reproduced from (A) for comparison (dashed lines). (**F**) Half-time of translocation was determined as in (B) and PKCβII-WT is reproduced from (B) for comparison. All data represent mean ± SEM. *** *P* < 0.001, **** *P* < 0.0001 by one-way ANOVA and Tukey post hoc test.

To assess if mutation of R42 is associated with higher basal activity, we co-overexpressed our FRET-based C-Kinase Activity Reporter (CKAR2; (56)) with mCherry-tagged WT PKCβII or disease-associated mutants R42H and R42P and monitored the change in FRET ratio following inhibitor addition (Figure 2C-D). Cells expressing CKAR2 alone (representing endogenous PKC activity, light blue trace) and those overexpressing mCherry-PKCβII-WT (dark blue trace) have low basal activity due to effective autoinhibition of the mature kinase. In cells expressing R42H or R42P mutants (red or grey traces, respectively), the activity drop following inhibitor addition was significantly greater than that observed for WT (Figure 2C), indicating higher basal activity. Thus, mutation of R42 to either His or Pro results in significantly higher basal signaling output, with R42P showing a larger increase, compared to the autoinhibited WT PKCβII (Figure 2D).

Given that both R42 and E655 mutations disrupted autoinhibition and are predicted to form an ion pair in our model, we assessed the effect of reversing the charge of both residues (R42E/E655R) on the translocation kinetics (Figure 2E-F). The rate of membrane-translocation of the double-mutant R42E/E655R was significantly faster than that of WT PKCβII and similar to that of R42E and E655R. Thus, charge reversal was not sufficient to rescue the autoinhibited conformation of PKCβII. This is unsurprising given the complex conformational transitions involved in the maturation of PKC. Mutation of the non-conserved (Figure 1B), neighboring E657 residue, which is not predicted to contribute to autoinhibition in our model, had no significant impact on translocation kinetics (Figure 2E-F). However, introducing an additional R42E mutation to the E657R mutant did increase translocation (green traces). These data show that perturbation of R42 or E655, but not adjacent E657, leads to a more open conformation of PKC with higher basal activity in a cellular context, and that charge reversal is insufficient to reclamp PKC into the tightly autoinhibited state.

### SCA14-associated R42P mutant evades PDBu-induced ubiquitination and degradation

The steady-state levels and activity of PKC are finely regulated by quality-control mechanisms that lead to the dephosphorylation, ubiquitination, and degradation of activated species (57–59). To examine the activation-induced degradation of the R42 mutants, we overexpressed YFP-tagged PKCβII WT or the indicated R42 mutants in COS7 cells and treated them with increasing concentrations of PDBu for 24 hours. Cells were lysed and relative PKC levels were determined by Western blot of whole-cell lysate (Figure 3A). PKCβII WT was dephosphorylated and degraded with increasing concentrations of PDBu as indicated by loss of the upper mobility, phosphorylated species (indicated by *; initial phosphorylation level shown in Figure 3B) and decrease in the total protein levels (Figure 3C). The highest concentration of PDBu (200 nM) resulted in degradation of WT and each mutant (Figure 3C), except for the R42P mutant, which was resistant to degradation. The initial levels of R42H, but not other mutants, were reduced compared to WT enzyme (approximately 90% relative to WT) (Figure 3A). This likely reflects the quality-control mechanisms which degrade PKC that is not properly autoinhibited and suggests that histidine at position 42 is particularly sensitive to downregulation. R42 mutants generally had decreased basal phosphorylation (Figure 3B), including R42P, consistent with a more open and phosphatase-sensitive conformation. Our results indicate that the R42P mutant is resistant to phorbol ester-induced downregulation.

**Figure 3.**
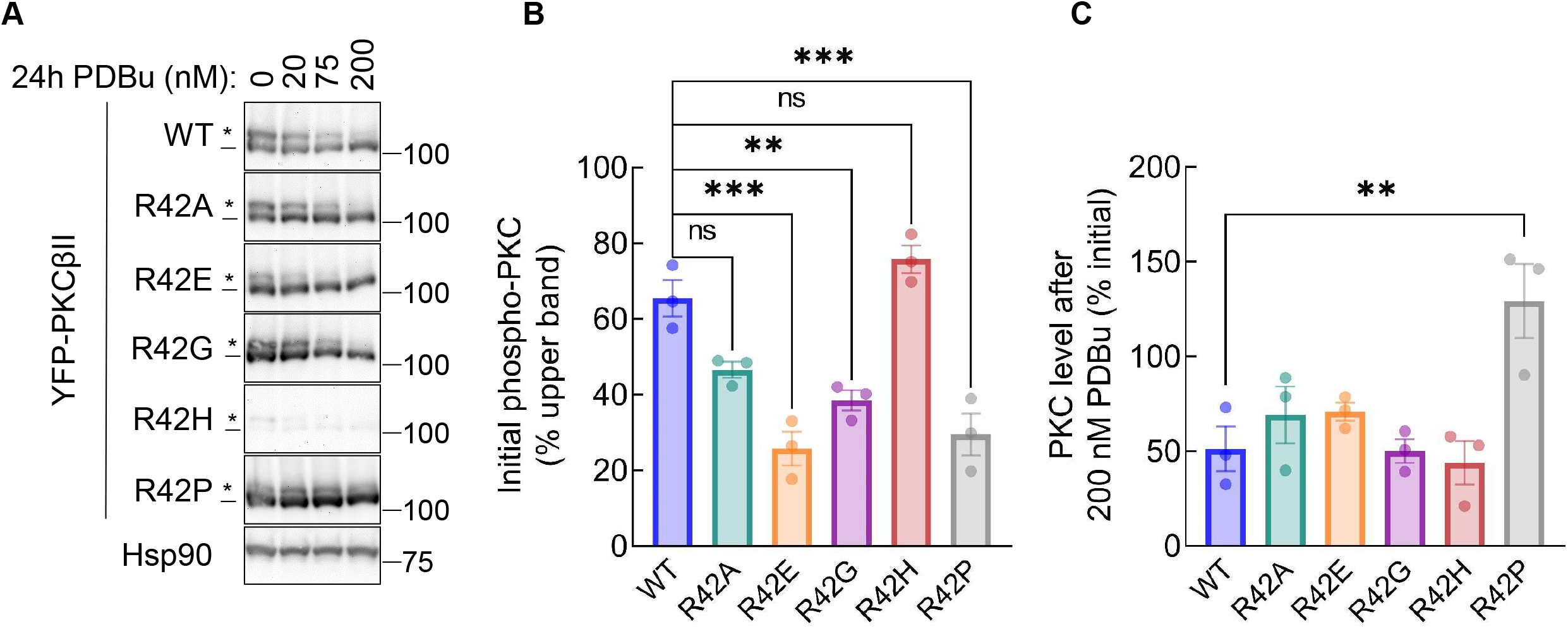
R42P SCA14-associated mutant is insensitive to phorbol ester-mediated downregulation. (**A**) Western blot of whole-cell lysate from COS7 cells transfected with YFP-tagged PKCβII wild-type or indicated R42 mutants. Cells were treated with indicated concentrations of PDBu for 24 hours before lysis. Blot is representative of three independent experiments. *, phosphorylated species; -, unphosphorylated species. (**B**) Quantification of percent phosphorylated PKC in DMSO-treated condition. (**C**) Quantification of percent change in PKC levels at 200 nM PDBu relative to 0 nM (DMSO) control for each transfection condition. Data represent mean ± SEM. ns = not significant, ** *P* < 0.01, *** *P* < 0.001 by one-way ANOVA and Tukey post-hoc test.

We next investigated the mechanism by which downregulation of PKCβII-R42P is impaired. COS7 cells transfected with YFP-tagged WT, R42H, or R42P PKCβII constructs for 48 hours were treated with the proteasome inhibitor MG-132 for 3 hours and then with PDBu for 30 minutes to allow ubiquitination, but not degradation, of PKC. PKC was immunoprecipitated from triton-soluble lysate and ubiquitination level of PKC was determined by Western blot (Figure 4A). The PDBu-mediated increase in ubiquitination was calculated by comparing the ubiquitination levels of untreated versus PDBu-treated sample for each protein (Figure 4B). Both WT and R42H showed significant increases in ubiquitination following phorbol ester treatment as seen previously (60). However, there was no increase in ubiquitination of the R42P mutant after stimulation with PDBu. These experiments reveal that the R42P mutant is resistant to PDBu-induced ubiquitination, providing a mechanism for its resistance to downregulation.

**Figure 4.**
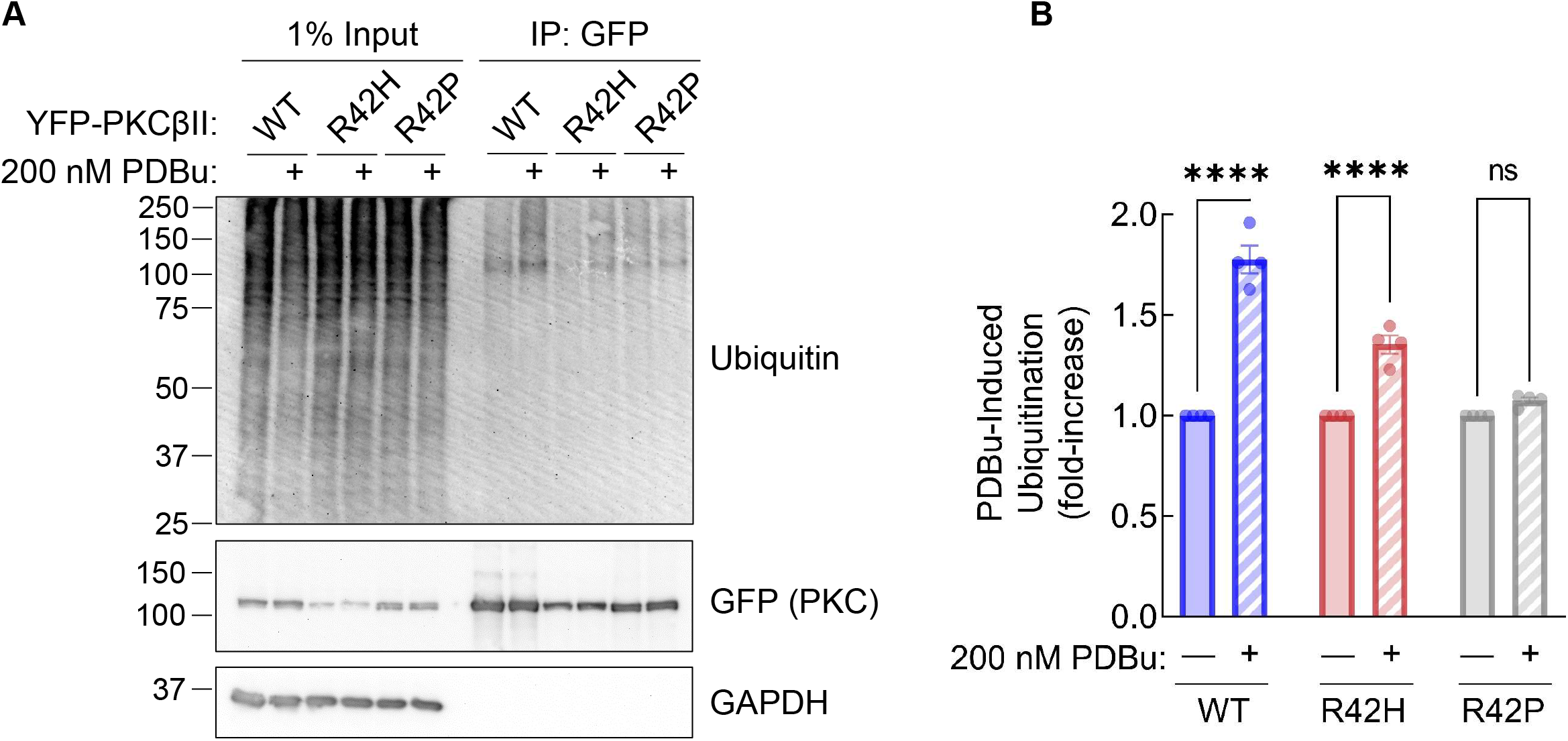
Ataxia-associated R42P mutation disrupts PDBu-induced ubiquitination of PKCβII. (**A**) Western blot of Triton-soluble lysate (left lanes) and YFP-PKCβII immunoprecipitated from COS7 cells using GFP-Trap^®^ Agarose. Cells were pre-treated with 20 μM MG-132 for 3 hours followed by 30 min of 200 nM PDBu treatment prior to lysis. Blots were probed with indicated antibodies. (**B**) Quantification of PDBu-induced ubiquitination of immunoprecipitated PKCβII. Relative ubiquitination was determined (Ubiquitin / PKC) for immunoprecipitated samples and each condition was normalized to DMSO-treated control (1.0) to determine fold-increase in ubiquitination after PDBu stimulation. Data represent mean ± SEM from four independent experiments. ns = not significant, **** *P* < 0.0001 by two-way ANOVA and Šídák’s multiple comparisons test.

### R42P mutant disrupts C1A domain stable residue network

Because R42P reduces autoinhibition while bypassing ubiquitination and degradation, an effect also observed upon deletion of the C1A domain (9), we hypothesize that a structural disruption of the C1A domain itself, rather than an interdomain effect, mediates its effect on PKC. We used molecular dynamics (MD) to explore how the domain responds to having an R42P substitution. For MD and analyses we used the C1A domain of PKCγ whose structure is solved (PDB: 2E73) and has 95% sequence identity to PKCβII; residue numbering as in PKCβII. Following 50 ns MD in triplicate, local spatial pattern (LSP) alignment-based protein residue networks (PRN) were created to detect stable regions and identify functional residues in the domain (61). Degree Centrality (DC) and Betweenness Centrality (BC) values for these networks were calculated as these measures have been shown to be effective in identifying functionally important nodes within a network (62, 63). As expected, two distinct “hubs” clustered around the two Zn^2+^ ions had high DC, indicating their local stability within the domain (Figure 5A, yellow and blue shading). To further examine residues with importance to domain connectivity, we created a scatterplot with DC and BC for each residue (Figure 5B). Residues with high BC represent key communicators, and while we found several Zn^2+^-binding residues (C50, C70, H75, C86) important for this global connectivity, the highest-scoring residue was V73 (green). This residue sits in the middle of the domain and appears to be a central node of the domain, connecting the two Zn^2+^ communities (Figure 5A, green residue). In contrast, R42 (blue residue) is on a flexible loop that extends outward and has low measures of DC and BC.

**Figure 5.**
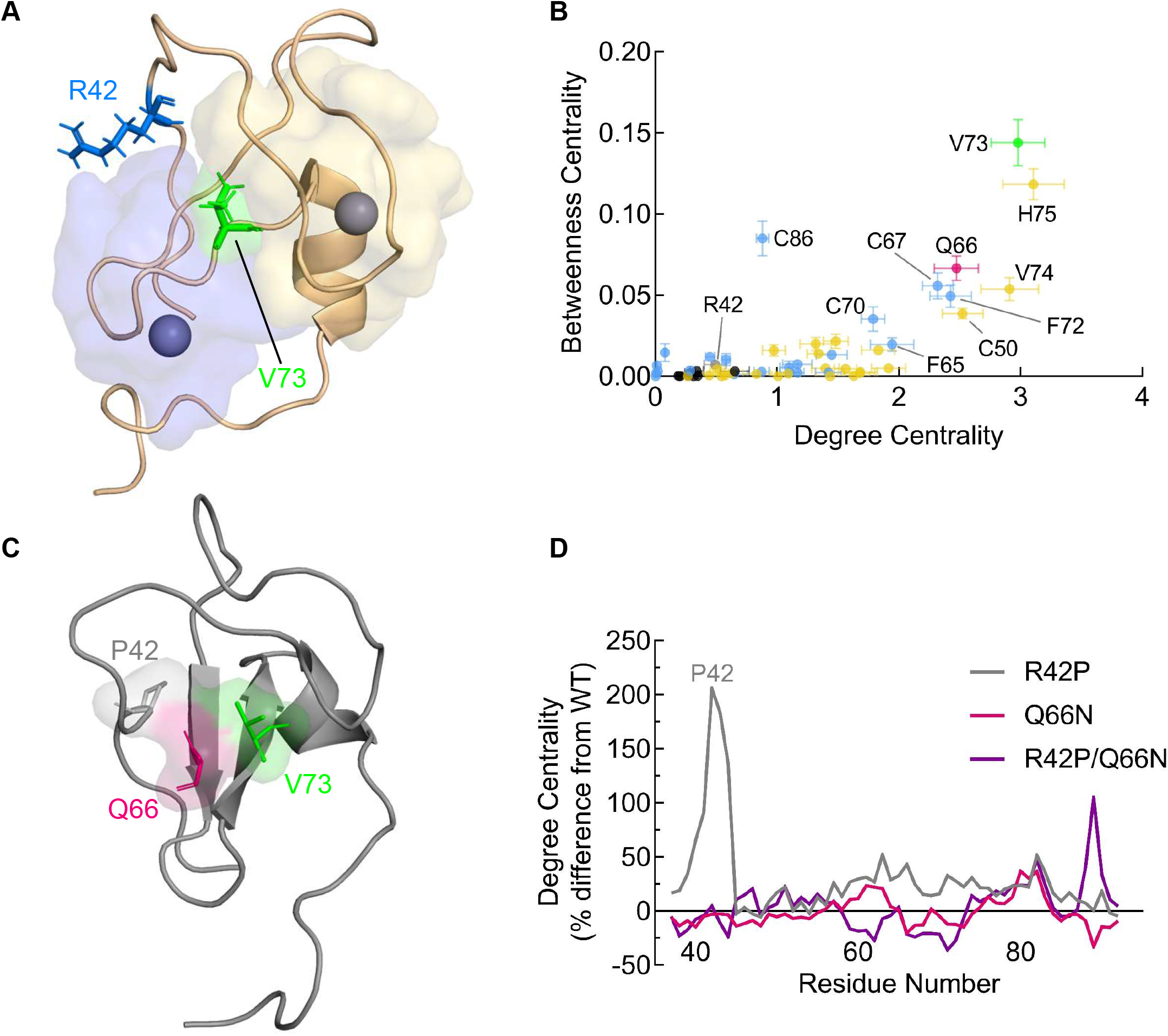
Molecular dynamics with local spatial pattern analysis reveals C1A domain organization and predicts disruption mechanism of R42P. (**A**) Structure of the PKCγ C1A domain (PDB: 2E73) shown with residues participating in hubs centered around Zn^2+^ ions colored in yellow and blue. Blue and green residues are R42 and V73, respectively. Note: residues numbered according to PKCβII sequence. (**B**) Molecular dynamics analysis of the PKCγ C1A domain (PDB: 2E73) was performed for 50 ns in triplicate followed by local spatial pattern alignment as described previously (61). Scatter plot represents the distribution of residues based on their degree and betweenness centralities. Standard errors calculated as described in Methods are shown and colored as in (A). (**C**) R42P modeled into C1A domain followed by molecular dynamics and LSP analysis shows potential interaction involving P42 (grey), Q66 (pink), and V73 (green). (**D**) Degree centrality of each residue in the C1A domain was determined from MD simulations of R42P (grey), Q66N (pink), and R42P/Q66N (purple) and the percent difference from WT of each residue is shown for each mutant.

Next, the MD and LSP was repeated with the R42P mutant modeled into the domain, and an interaction was seen involving P42, Q66, and V73 in multiple frames of the MD (Figure 5C; residues in grey, pink, and green, respectively). Strikingly, the DC of each residue on the loop containing P42 increased, indicating stabilization of the loop resulting in a denser group of residues with this mutation present (Figure 5D, grey trace). Because the hydrophobic side chain of the proline is predicted to bind to the hydrophobic core of the domain through the Q66 side chain in the MD simulation, we hypothesized that shortening this residue to an asparagine would prevent this interaction, rescue loop flexibility, and repair domain dynamics. In line with this, the double-mutant R42P/Q66N showed restoration of DC, especially surrounding the loop containing P42 (Figure 5D, purple trace). Mutation of Q66 alone did not lead to substantial changes in DC (Figure 5D, pink trace). If altered dynamics of the C1A domain are resulting in impaired degradation of the R42P mutant, these results suggest a strategy to restore C1A function of the R42P mutant would be to introduce a Q66N mutation, which our results indicate will create stable communities within the domain similar to the WT.

### Degradation of double-mutant R42P/Q66N is restored to WT levels as predicted by MD and LSP

Our MD results indicate that R42P perturbs C1A domain dynamics through introducing structural rigidity and additional interactions within the domain, including with Q66. To test our *in silico* prediction that mutating Q66 to asparagine in the R42P mutant may restore normal C1A function, we overexpressed YFP-tagged PKCβII WT, R42P, Q66N, and R42P/Q66N in COS7 cells, stimulated them with PDBu for 24 hours, and analyzed PKC levels by Western blot (Figure 6A). As previously observed, the R42P mutant was not degraded even at this high PDBu concentration (Figure 6B, grey bar). Q66N also had significantly reduced degradation compared to WT (P = 0.0068) suggesting that mutation of this residue, which our initial MD/LSP revealed has relatively high DC and BC, can also alter C1A function (Figure 5B, pink point and Figure 6B, pink bar). Mutation of both R42P and Q66N together resulted in degradation of PKCβII after 24 hours of PDBu treatment. Furthermore, the degree of degradation was the same as WT (Figure 6B, purple bar). These results suggest that the double-mutant has restored C1A domain-mediated downregulation of PKCβII. Therefore, applying MD and LSP alignment methods allowed us to determine and repair a mechanism of C1A domain disruption which is blocking the ubiquitination and degradation of PKC.

**Figure 6.**
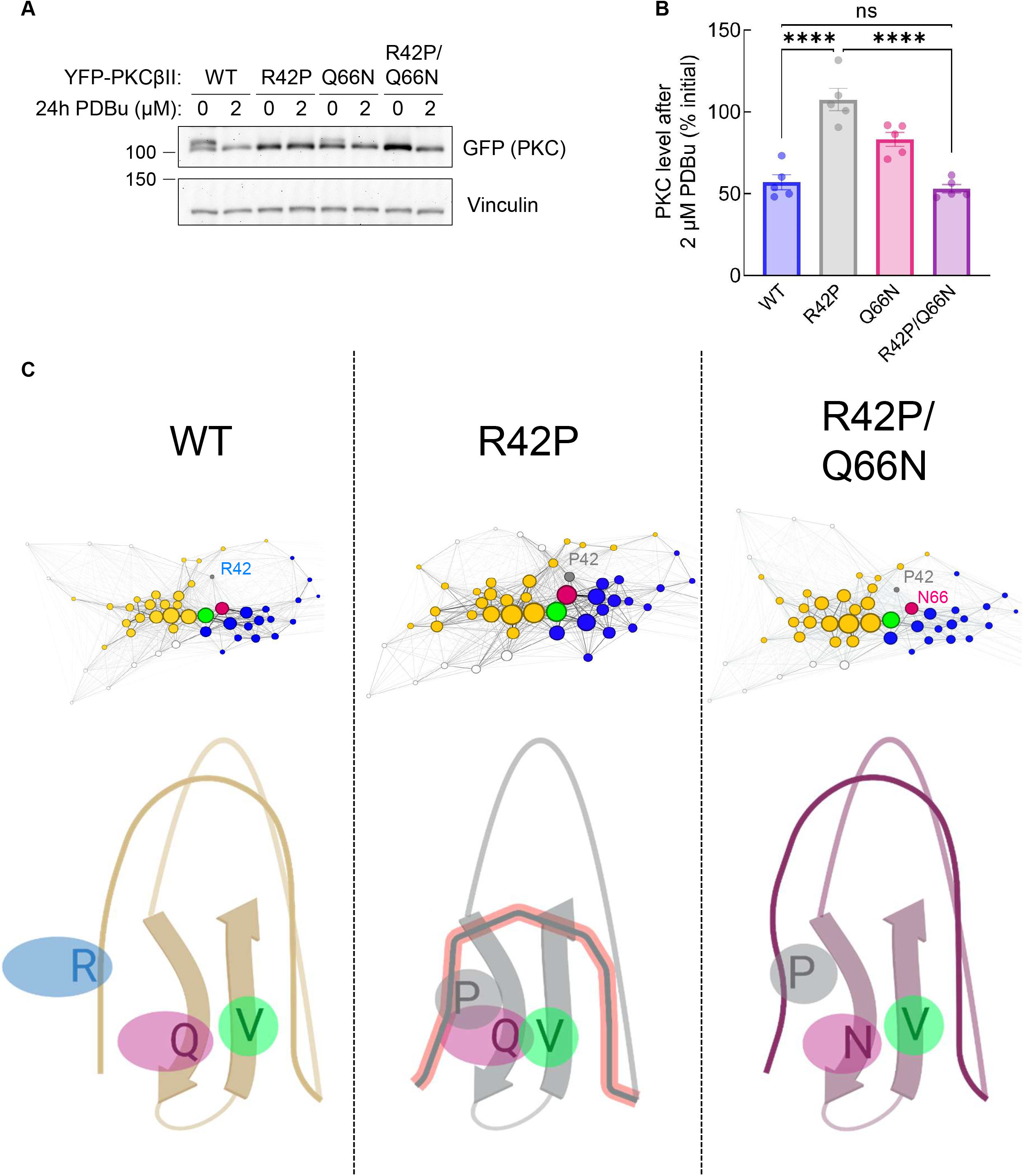
Q66N mutation restores degradation of R42P-mutated PKCβII. (**A**) Western blot of whole-cell lysate from COS7 cells transfected with YFP-tagged PKCβII WT, R42P, Q66N, or the double-mutant R42P/Q66N. Cells were treated with DMSO or 2 μM PDBu for 24 hours before lysis. Blot is representative of five independent experiments. (**B**) Quantification of (A) showing percent change in PKC levels at 2 μM PDBu relative to DMSO control. ns = not significant, **** P < 0.0001 by one-way ANOVA and Tukey post-hoc test. (**C**) Proposed mechanism of C1A disruption by R42P mutation (middle panel) and restoration of domain dynamics through double mutant R42P/Q66N (right panel). Maps represent LSP data visualized using Gephi software. Each node represents an amino acid with the diameter corresponding to Degree Centrality. Yellow and blue nodes indicate hubs of residues participating in Zn^2+^ binding as in Figure 5A. Grey, pink, and green nodes are R42 or P42, Q66 or N66, and V73, respectively. Figure created using BioRender.com.

## Discussion

Mounting evidence points to cancer-associated variants in PKC generally being loss-of-function and variants present in neurodegenerative disease having gain-of-function alterations (39, 64, 65). In this study, we show that mutation of a conserved residue in the C1A domain of cPKC leads to opposing phenotypes depending on the disease context. Mutation of R42 to histidine (cancer-associated) or proline (SCA14-associated) similarly impaired autoinhibition (Figure 2). However, whereas mutation to histidine triggered quality-control degradation, mutation to proline altered the fold of the C1A domain to protect PKC from activator-induced ubiquitination and degradation (Figures 3 and 4). MD revealed that the stable residue networks of the C1A domain are disrupted by R42P mutation (Figure 5). Importantly, we restored sensitivity to downregulation of this mutant by with a compensating mutation predicted from LSP analysis (Figures 6). Our data reveal how complex regulatory mechanisms control PKC structure and function such that mutation of the same residue can have opposing outcomes in disease.

The predicted ion pair between R42 in the C1A domain and E655 in the regulatory C-tail is conserved throughout evolution, suggesting a key role in maintenance of normal PKC function (Figure 1B). Consistent with this, we found that mutation of the invariant R42 to any residue tested and mutation of E655 to arginine resulted in disrupted autoinhibition of PKC. These results point to a key functional role of this ion pair in maintaining effective autoinhibition of cPKC isozymes. Thus, it is unsurprising that the ion pair is targeted in disease as a way to impair PKC function. Not only is R42 mutated to histidine in two separate cancers, but E655 mutation to tyrosine, lysine, histidine, and aspartate has also been identified in cancers across the cPKC family (42).

A striking finding from our analysis was that mutation of R42 to proline prevented phorbol ester-induced ubiquitination and downregulation, rendering the PKC insensitive to quality-control mechanisms (32, 33, 60). To understand the structural basis of R42P disruption of PKC function, we used molecular dynamics simulations followed by LSP alignment to study changes in stable regions of the C1A domain. This approach has been shown to out-perform traditional interaction-based methods to identify critical regulatory residues in protein kinase A (61). Applying this method, we observed a rewiring of the C1A domain when the R42P mutation was introduced (Figure 5C-D). The proline is predicted to form a hydrophobic network with Q66 and V73, the latter of which is a key node of the domain that connects the two Zn^2+^-binding hubs. This interaction also collapses a ligand-binding loop leading to dramatic changes in the stable residue network of the domain (Figure 6C, middle panel). From the LSP analysis, we predicted that shortening of the hydrophobic sidechain of Q66 to an asparagine would release V73 from this hydrophobic interaction to restore hub communication and domain centralities. Double-mutation of R42P and Q66N repaired loop flexibility and degree centrality of most residues in the domain back to that of WT (Figure 5D, Figure 6C, right panel). This double-mutant (R42P/Q66N) was degraded in response to phorbol esters, suggesting that stable PRNs in the C1A domain are critical to agonist-induced degradation of PKC. These results provide a framework for studying the effect of disease-associated mutations and generate testable hypotheses based on PRN maps to restore dynamics and communication within protein domains.

R42A mutation in PKCα was previously shown to have higher *in vitro* basal activity than WT enzyme (51), and this mutant also translocated faster to the plasma membrane (Figure 2A-B). Mutation of the neighboring F43 also increases membrane translocation rate, and mutation of this residue in PKCδ (F165) leads to a more open conformation, indicating that this region of the C1A domain contributes to autoinhibition of PKC isozymes (53). C1A-domain synthesized peptides with the R42P mutation (R41P in PKCγ) do not bind PDBu (66); however, the full-length protein is able to readily translocate to the plasma membrane in response to stimulation in cells. Phorbol ester-induced translocation has previously been shown to be mediated primarily by the C1B domain binding phorbol esters (23, 27). Thus, although both the C1A and C1B domains bind phorbol esters as isolated domains (23, 67), in the context of the full-length mature enzyme, the C1B is the diacylglycerol/phorbol ester sensor. Therefore, the increased rate of translocation observed with the R42P mutant reflects a more open PKC with a more accessible C1B domain (68).

The resistance of the R42P mutant to ubiquitination suggests a key role of the C1A domain in either being ubiquitinated or providing a docking site for an E3 ubiquitin ligase. The E3 ubiquitin ligase MDM2 is known to ubiquitinate phorbol-ester activated PKCβII (60, 69). The PKCβII—MDM2 interaction was shown to be mediated by the C1 domains of PKC, so the structural disruption of the C1A caused by R42P mutation may disrupt this interaction. SCA14 mutants tend to cluster in the C1 domains of PKCγ (9, 37, 43, 66), so there may be a common mechanism involving disruption of the MDM2 binding site or interface which would block ubiquitination and downregulation of activated PKC. Biochemical characterization of other SCA14 mutants in PKCγ have shown similar defects in phorbol ester-mediated downregulation while also having increased basal activity (9). Importantly deletion of the entire C1A domain also shares this phenotype. The role of the C1A domain in mediating the downregulation of PKC remains to be elucidated.

In summary, our work adds to a growing field of evidence pointing towards PKC as a tightly regulated signaling node that is perturbed in disease. Consistent with previous studies that examined PKC in the context of cancer and neurodegenerative diseases, we found that these mechanisms of loss- vs gain-of-function, respectively, hold true even for mutations at the same residue. Applying both biochemical and structural techniques allowed us to repair the downregulation of a disease-associated mutant and gain new insights into the function of specific residues in the C1A domain.

## Methods

### Plasmid Constructs, Antibodies, and Reagents

The CKAR2 (56) and mpCFP (54) were described previously. Human PKCβII was YFP-tagged at the N-terminus in a pcDNA3 vector using Gateway Cloning (Life Technologies) as described in (6). All mutants were generated using QuikChange site-directed mutagenesis (Agilent) following the manufacturer’s instructions. Antibodies against GFP (catalog no. 2555S), Vinculin (catalog no. 4650S), Ubiquitin (catalog no. 3933S), and GAPDH (14C10, catalog no. 2118) were from Cell Signaling Technologies and used at 1:1000 dilution. Hsp90 antibody was from BD Biosciences (catalog no. 610419) used at 1:1000 dilution. HRP-conjugated anti-rabbit (catalog no. 401315) and anti-mouse (catalog no. 401215) secondary antibodies and BSA (catalog no. 12659) were from Millipore. All antibodies were diluted in 1% BSA dissolved in PBS-T (1.5 mM Sodium Phosphate Monobasic, 8 mM Sodium Phosphate Dibasic, 150 mM NaCl, 0.05% Tween-20) with 0.25 mM thimerosal (Thermo Scientific, catalog no. J61799.14). PDBu (catalog no. 524390) and MG-132 (catalog no. 474790) were purchased from Calbiochem. Gö6983 (catalog no. 285) was purchased from Tocris. Bradford reagent (catalog no. 500-0006), protein standards ladder (catalog no. 161-0394), bis/acrylamide solution (catalog no. 161-0156), and polyvinylidene difluoride (PVDF) (catalog no. 162-0177) were purchased from Bio-Rad. Luminol (catalog no. A-8511) and p-coumaric acid (catalog no. C-9008) used to make chemiluminescent substrate solution were purchased from Sigma-Aldrich.

### Cell Lysis and Western Blotting

Prior to lysis, cells were washed with Dulbecco’s phosphate-buffered saline (DPBS) (Corning, catalog no. 21-031-CV). Cells were lysed in Phosphate Lysis Buffer pH 7.4 containing 50 mM sodium phosphate (38 mM sodium phosphate dibasic, 12 mM sodium phosphate monobasic), 1 mM sodium pyrophosphate, 20 mM sodium fluoride, 2 mM EDTA, and 1% Triton X-100. Lysis buffer was supplemented with 1 mM phenylmethylsulfonyl fluoride (PMSF), 50 μg/ml leupeptin, 1 mM Na3VO4, 2 mM benzamidine, 1 μM microcystin, and 1 mM DTT added immediately prior to lysis. Lysates were collected by scraping and whole-cell lysates were briefly sonicated prior to quantification by Bradford Assay. Samples were boiled in sample buffer containing 250 mM Tris HCl, 8% (w/v) SDS, 40% (v/v) glycerol, 80 μg/ml bromophenol blue, and 2.86 M β-mercaptoethanol for 5 minutes at 95 °C. Unless otherwise noted, 20 μg protein per sample was analyzed by SDS-PAGE using 6% acrylamide gels to visualize phosphorylation-induced mobility shifts. Gels were transferred to membranes (PVDF) for three hours at 80 V in transfer buffer (200 mM Glycine, 25 mM Tris Base, 20% Methanol) at 4 °C. Membranes were blocked in 5% milk dissolved in PBS-T for 30 minutes at room temperature then washed with PBS-T for 5 minutes three times before incubating with primary antibody overnight at 4 °C with rocking. Membranes were washed for 5 minutes three times in PBS-T, secondary antibodies were added for 1 hour at room temperature, and the wash step was repeated before developing with chemiluminescence. Chemiluminescent solution (100 mM Tris pH 8.5, 1.25 mM luminol, 198 μM coumaric acid, and 1% H_2_O_2_) was added to membranes for 2 minutes then imaged on a FluorChem Q imaging system (ProteinSimple).

### Cell Culture and Transfection

COS7 cells were maintained in Dulbecco’s modified Eagle’s medium (Corning, catalog no. 10-013-CV) containing 10% fetal bovine serum (Atlanta Biologicals, catalog no. S11150) and 1% penicillin/streptomycin (Gibco, catalog no. 15-140-122) at 37°C in 5% CO2. Cells were periodically tested for Mycoplasma contamination by a PCR-based method (70). Transient transfections were carried out using a Lipofectamine 3000 kit (Thermo Fisher Scientific) per the manufacturer’s instructions, and constructs were allowed to express for 24 hours prior to drug treatment or imaging experiments.

### FRET Imaging and Analysis

2 x 10^5^ COS7 cells were seeded into plates (Corning, catalog no. 430165) containing glass cover slips (Fisherbrand, catalog no. 12545102) glued on using SYLGARD 184 Silicone Elastomer Kit (Dow, catalog no. 04019862) and cells were transfected 24 hours after seeding. For CKAR assays, cells were co-transfected with 1 μg mCherry-PKCβII constructs and 1 μg CKAR2 DNA (56). For translocation assays, cells were cotransfected with 800 ng YFP-PKCβII and 400 ng MyrPalm-CFP (54). 24 hours posttransfection, cells were imaged in 2 mL Hank’s Balanced Salt Solution (Corning, catalog no. 21-022-CV) with 1 mM CaCl2 added fresh prior to imaging. Images were acquired on a Zeiss Axiovert 200M microscope (Carl Zeiss Micro-Imaging Inc.) using an Andor iXonUltra 888 digital camera (Oxford Instruments) controlled by MetaFluor software (Molecular Devices) version 7.10.1.161. Filter sets and parameters for imaging are described in (71). Background signal was subtracted for each wavelength from area containing no cells. One region per cell was selected for quantification by tracing an area excluding the nucleus for CKAR imaging and containing the whole cell for translocation. Images were acquired every 15 seconds, and baseline images were acquired for 3 minutes. Agonist (200 nM PDBu) or inhibitor (1 μM Gö6983) was added dropwise to the dish in-between acquisitions. For CKAR activity assays, FRET ratios for each cell were normalized to the 15-minute end point and basal activity was calculated by averaging the initial normalized FRET ratio. For translocation assays, half-times were calculated by fitting the data to a non-linear regression using a plateau followed by one-phase association equation with X0 = 3 min. Data were normalized to the maximum FRET ratio for each cell (100%) to calculate percent translocation. Data represent mean ± SEM for cells from at least three independent experiments.

### Phorbol Ester Downregulation Assay

COS7 cells were seeded into six-well plates at 2 x 10^5^ cells per well. Cells were transfected after 24 hours with 500 ng of indicated constructs and allowed to incubate for 24 hours. For dose-response experiments, cells were treated with up to 2 μM PDBu or control (DMSO) for 24 hours prior to lysis.

### Ubiquitination Assay

COS7 cells were seeded into 10 cm plates 24 hours prior to transfection with 2.5 μg DNA per plate. After 48 hours of transfection, cells were treated with 20 μM MG-132 for 3 hours followed by 200 nM PDBu for 30 minutes. Cells were lysed in Phosphate Lysis Buffer (same recipe as above) and clarified by centrifugation for 10 min at 12,000 x g. Triton-soluble lysate was quantified using Bradford Assay and 500 μg was used for IP. GFP-Trap® Agarose or control agarose (Proteintech, product code: gta for GFP-Trap® and bab for binding control) was added and incubated at 4 °C overnight on a rocker with gentle motion. Beads were sedimented by centrifugation at 2,500 x g for 2 min and supernatant was removed. Four washes of the agarose in Phosphate Lysis Buffer were performed with centrifugation in-between each. After the final wash, sample buffer was added, and the agarose was boiled for 5 min at 95 °C for elution. Samples were centrifuged and the supernatant was saved for analysis by SDS-PAGE and Western blot.

### Phylogenetic Analysis

cPKC homologs were gathered using BlastP against the NCBI NRAA database, and other classes of PKC were removed by BlastP against a small database of diverse PKC isozymes. Sequences were aligned in Clustal Omega (72) and edited in JalView (73). The sequence logo was created with a set of representative sequences: one per vertebrate class for PKCα, PKCβ and PKCγ, and a diverse set of invertebrate bilaterian PKC isozymes, one per phylum. The sequence logo was visualized by Weblogo (http://weblogo.berkeley.edu) (74).

### Molecular Dynamics and Local Spatial Pattern alignment

The model for simulations was based on the structure of the C1A domain of PKC (PDB: 2E73). The Protein Preparation Wizard was used to model the charge states of ionizable residues at neutral pH, and hydrogens and counter ions were added. The resulting model was solvated in a cubic box of TIP4P-EW and 150 mM KCl with a 10 Å buffer in AMBER tools. Energy minimization, heating, and equilibration steps were performed using AMBER16 (75). Systems were first minimized by 900 steps of hydrogen-only minimization, 2000 steps of solvent minimization, 2000 steps of sidechain minimization, and 5000 steps of all-atom minimization. The systems were then heated from 0°K to 100°K linearly over 250 ps with 2 fs time-steps and 5.0 kcal·mol·Å position restraints on the protein backbone under Constant Volume. Langevin dynamics were used to control the temperature using a collision frequency of 1.0 ps-1. Then the systems were heated from 100°K to 300°K linearly over 200 ps with 2 fs time-steps and 5.0 kcal·mol·Å position restraints on the protein backbone under Constant pressure. Constant pressure equilibration was performed with a 10 Å non-bonded cut-off for 250 ps, with 5.0 kcal·mol·Å position protein backbone restrained, followed by 250 ps of unrestrained equilibration. Finally, production simulations were conducted on a GPU-enabled AMBER16 in triplicate for each construct for 50 ns (76).

To generate LSP-based networks, three 50 ns MD trajectories were split into 15 intervals 10 ns each. From each interval, 100 structures were extracted with a 0.1 ns step, and LSP-alignment was performed within each group of 100 structures using inhouse software in all-to-all manner as described earlier (61). Briefly, prior to alignment, each protein structure was represented as an undirected weighted graph with residues as nodes and links were formed between residues if the distance between the corresponding Cα atoms was less than 12Å. Each link was described by four numbers that represented the orientation of the CαCβ vectors of the two residues: three distances (Cα1-Cα2, Cα1-Cβ2, Cα2-Cβ1) and the dihedral angle θ (Cβ1-Cα1-Cα2-Cβ2). The result of each LSP-alignment was represented as a graph where links were created between two residues if their orientation was similar in the two structures being compared. Similar orientations were defined by preset cut-off levels for the three distances and the angle: ΔCα1-Cα2 < 0.2Å, ΔCα1-Cβ2 <0.45Å, ΔCα2-Cβ1<0.45Å, Δθ<10°. The all-to-all LSP-alignment of 100 structures within each 10 ns interval produced 4950 adjacency matrices, which were then averaged and normalized Degree and Betweenness centrality were calculated using igraph R library (version 1.2.5) (77). To calculate betweenness centrality weights (W) were converted to distances (D) using the formula: D=-log(W). The 15 values of degree and betweenness centrality were averaged, and standard errors were calculated.

### Quantification and Statistical Analysis

For imaging experiments, intensity values and FRET ratios were acquired using MetaFluor software and normalized as described above. Western blots were quantified by densitometry using AlphaView (ProteinSimple) version 3.4.0.0. Statistical tests described in figure legends were performed using Prism (GraphPad Software) version 9.5.0. Structures were modeled using PyMOL version 2.3.0 (Schrödinger, LLC).

## Abbreviations

PKC: protein kinase C
cPKC: conventional protein kinase C
AD: Alzheimer’s Disease
SCA14: Spinocerebellar Ataxia Type-14
DG: diacylglycerol
PIP2: phosphatidylinositol-4,5-bisphosphate
PDK-1: phosphoinositide-dependent kinase 1
YFP: yellow fluorescent protein
CFP: cyan fluorescent protein
PDBu: phorbol 12,13-dibutryate
FRET: fluorescence resonance energy transfer
CKAR: C kinase activity reporter
MD: molecular dynamics
LSP: local spatial pattern alignment
PRN: protein residue network
DC: degree centrality
BC: betweenness centrality

## Data Availability

All supporting data are included within the main article.

## Acknowledgements

We thank members of the Newton and Taylor laboratories for their many helpful discussions and edits to the manuscript.

## References

1. Nishizuka Y. Protein kinase C and lipid signaling for sustained cellular responses. FASEB journal: official publication of the Federation of American Societies for Experimental Biology. 1995;9(7):484–96.

2. Newton AC. Protein kinase C: perfectly balanced. Critical reviews in biochemistry and molecular biology. 2018;53(2):208–30.

3. Kajimoto T, Caliman AD, Tobias IS, Okada T, Pilo CA, Van AN, et al. Activation of atypical protein kinase C by sphingosine 1-phosphate revealed by an aPKC-specific activity reporter. Science signaling. 2019;12(562).

4. Newton AC. Protein kinase C: structural and spatial regulation by phosphorylation, cofactors, and macromolecular interactions. Chemical reviews. 2001;101(8):2353–64.

5. Callender JA, Newton AC. Conventional protein kinase C in the brain: 40 years later. Neuronal signaling. 2017;1(2):Ns20160005.

6. Antal CE, Hudson AM, Kang E, Zanca C, Wirth C, Stephenson NL, et al. Cancer-associated protein kinase C mutations reveal kinase’s role as tumor suppressor. Cell. 2015;160(3):489–502.

7. Alfonso SI, Callender JA, Hooli B, Antal CE, Mullin K, Sherman MA, et al. Gain-of-function mutations in protein kinase Cα (PKCα) may promote synaptic defects in Alzheimer’s disease. Science signaling. 2016;9(427):ra47.

8. Callender JA, Yang Y, Lordén G, Stephenson NL, Jones AC, Brognard J, et al. Protein kinase Cα gain-of-function variant in Alzheimer’s disease displays enhanced catalysis by a mechanism that evades down-regulation. Proceedings of the National Academy of Sciences of the United States of America. 2018;115(24):E5497–e505.

9. Pilo CA, Baffi TR, Kornev AP, Kunkel MT, Malfavon M, Chen DH, et al. Mutations in protein kinase Cγ promote spinocerebellar ataxia type 14 by impairing kinase autoinhibition. Science signaling. 2022;15(753):eabk1147.

10. Abrahamsen H, O’Neill AK, Kannan N, Kruse N, Taylor SS, Jennings PA, et al. Peptidyl-prolyl isomerase Pin1 controls down-regulation of conventional protein kinase C isozymes. The Journal of biological chemistry. 2012;287(16):13262–78.

11. Baffi TR, Van AN, Zhao W, Mills GB, Newton AC. Protein Kinase C Quality Control by Phosphatase PHLPP1 Unveils Loss-of-Function Mechanism in Cancer. Molecular cell. 2019;74(2):378–92.e5.

12. Dutil EM, Toker A, Newton AC. Regulation of conventional protein kinase C isozymes by phosphoinositide-dependent kinase 1 (PDK-1). Current biology: CB. 1998;8(25):1366–75.

13. Facchinetti V, Ouyang W, Wei H, Soto N, Lazorchak A, Gould C, et al. The mammalian target of rapamycin complex 2 controls folding and stability of Akt and protein kinase C. The EMBO journal. 2008;27(14):1932–43.

14. Behn-Krappa A, Newton AC. The hydrophobic phosphorylation motif of conventional protein kinase C is regulated by autophosphorylation. Current biology: CB. 1999;9(14):728–37.

15. Baffi TR, Newton AC. Protein kinase C: release from quarantine by mTORC2. Trends in biochemical sciences. 2022;47(6):518–30.

16. Baffi TR, Lordén G, Wozniak JM, Feichtner A, Yeung W, Kornev AP, et al. mTORC2 controls the activity of PKC and Akt by phosphorylating a conserved TOR interaction motif. Science signaling. 2021;14(678).

17. Keranen LM, Dutil EM, Newton AC. Protein kinase C is regulated in vivo by three functionally distinct phosphorylations. Current biology: CB. 1995;5(12):1394–403.

18. Orr JW, Newton AC. Requirement for negative charge on “activation loop” of protein kinase C. The Journal of biological chemistry. 1994;269(44):27715–8.

19. Edwards AS, Faux MC, Scott JD, Newton AC. Carboxyl-terminal phosphorylation regulates the function and subcellular localization of protein kinase C betaII. The Journal of biological chemistry. 1999;274(10):6461–8.

20. Biondi RM, Cheung PC, Casamayor A, Deak M, Currie RA, Alessi DR. Identification of a pocket in the PDK1 kinase domain that interacts with PIF and the C-terminal residues of PKA. The EMBO journal. 2000;19(5):979–88.

21. Yang J, Cron P, Thompson V, Good VM, Hess D, Hemmings BA, et al. Molecular mechanism for the regulation of protein kinase B/Akt by hydrophobic motif phosphorylation. Molecular cell. 2002;9(6):1227–40.

22. Hurley JH, Newton AC, Parker PJ, Blumberg PM, Nishizuka Y. Taxonomy and function of C1 protein kinase C homology domains. Protein science: a publication of the Protein Society. 1997;6(2):477–80.

23. Giorgione J, Hysell M, Harvey DF, Newton AC. Contribution of the C1A and C1B domains to the membrane interaction of protein kinase C. Biochemistry. 2003;42(38):11194–202.

24. Das J, Rahman GM. C1 domains: structure and ligand-binding properties. Chemical reviews. 2014;114(24):12108–31.

25. Katti SS, Krieger IV, Ann J, Lee J, Sacchettini JC, Igumenova TI. Structural anatomy of Protein Kinase C C1 domain interactions with diacylglycerol and other agonists. Nature communications. 2022;13(1):2695.

26. Sommese RF, Ritt M, Swanson CJ, Sivaramakrishnan S. The Role of Regulatory Domains in Maintaining Autoinhibition in the Multidomain Kinase PKCα. The Journal of biological chemistry. 2017;292(7):2873–80.

27. Antal CE, Violin JD, Kunkel MT, Skovsø S, Newton AC. Intramolecular conformational changes optimize protein kinase C signaling. Chemistry & biology. 2014;21(4):459–69.

28. House C, Kemp BE. Protein kinase C contains a pseudosubstrate prototope in its regulatory domain. Science (New York, NY). 1987;238(4834):1726–8.

29. Berridge MJ. Inositol trisphosphate and diacylglycerol: two interacting second messengers. Annual review of biochemistry. 1987;56:159–93.

30. Evans JH, Murray D, Leslie CC, Falke JJ. Specific translocation of protein kinase Calpha to the plasma membrane requires both Ca2+ and PIP2 recognition by its C2 domain. Molecular biology of the cell. 2006;17(1):56–66.

31. Johnson JE, Giorgione J, Newton AC. The C1 and C2 domains of protein kinase C are independent membrane targeting modules, with specificity for phosphatidylserine conferred by the C1 domain. Biochemistry. 2000;39(37):11360–9.

32. Van AN, Kunkel MT, Baffi TR, Lordén G, Antal CE, Banerjee S, et al. Protein kinase C fusion proteins are paradoxically loss-of-function in cancer. The Journal of biological chemistry. 2021:100445.

33. Hansra G, Garcia-Paramio P, Prevostel C, Whelan RD, Bornancin F, Parker PJ. Multisite dephosphorylation and desensitization of conventional protein kinase C isotypes. The Biochemical journal. 1999;342 ( Pt 2)(Pt 2):337–44.

34. Gao T, Brognard J, Newton AC. The phosphatase PHLPP controls the cellular levels of protein kinase C. The Journal of biological chemistry. 2008;283(10):6300–11.

35. Hsu AH, Lum MA, Shim KS, Frederick PJ, Morrison CD, Chen B, et al. Crosstalk between PKCα and PI3K/AKT Signaling Is Tumor Suppressive in the Endometrium. Cell reports. 2018;24(3):655–69.

36. Lordén G, Wozniak JM, Doré K, Dozier LE, Cates-Gatto C, Patrick GN, et al. Enhanced activity of Alzheimer disease-associated variant of protein kinase Cα drives cognitive decline in a mouse model. Nature communications. 2022;13(1):7200.

37. Chen DH, Brkanac Z, Verlinde CL, Tan XJ, Bylenok L, Nochlin D, et al. Missense mutations in the regulatory domain of PKC gamma: a new mechanism for dominant nonepisodic cerebellar ataxia. American journal of human genetics. 2003;72(4):839–49.

38. Sun YM, Lu C, Wu ZY. Spinocerebellar ataxia: relationship between phenotype and genotype - a review. Clinical genetics. 2016;90(4):305–14.

39. Pilo CA, Newton AC. Two Sides of the Same Coin: Protein Kinase C γ in Cancer and Neurodegeneration. Frontiers in cell and developmental biology. 2022;10:929510.

40. Zhang G, Kazanietz MG, Blumberg PM, Hurley JH. Crystal structure of the cys2 activator-binding domain of protein kinase C delta in complex with phorbol ester. Cell. 1995;81(6):917–24.

41. Jones AC, Taylor SS, Newton AC, Kornev AP. Hypothesis: Unifying model of domain architecture for conventional and novel protein kinase C isozymes. IUBMB life. 2020;72(12):2584–90.

42. Cerami E, Gao J, Dogrusoz U, Gross BE, Sumer SO, Aksoy BA, et al. The cBio cancer genomics portal: an open platform for exploring multidimensional cancer genomics data. Cancer discovery. 2012;2(5):401–4.

43. Chen DH, Cimino PJ, Ranum LP, Zoghbi HY, Yabe I, Schut L, et al. The clinical and genetic spectrum of spinocerebellar ataxia 14. Neurology. 2005;64(7):1258–60.

44. Parissenti AM, Kirwan AF, Kim SA, Colantonio CM, Schimmer BP. Inhibitory properties of the regulatory domains of human protein kinase Calpha and mouse protein kinase Cepsilon. The Journal of biological chemistry. 1998;273(15):8940–5.

45. Lopez-Garcia LA, Schulze JO, Fröhner W, Zhang H, Süss E, Weber N, et al. Allosteric regulation of protein kinase PKCζ by the N-terminal C1 domain and small compounds to the PIF-pocket. Chemistry & biology. 2011;18(11):1463–73.

46. Kirwan AF, Bibby AC, Mvilongo T, Riedel H, Burke T, Millis SZ, et al. Inhibition of protein kinase C catalytic activity by additional regions within the human protein kinase Calpha-regulatory domain lying outside of the pseudosubstrate sequence. The Biochemical journal. 2003;373(Pt 2):571–81.

47. Slater SJ, Seiz JL, Cook AC, Buzas CJ, Malinowski SA, Kershner JL, et al. Regulation of PKC alpha activity by C1-C2 domain interactions. The Journal of biological chemistry. 2002;277(18):15277–85.

48. Hommel U, Zurini M, Luyten M. Solution structure of a cysteine rich domain of rat protein kinase C. Nature structural biology. 1994;1(6):383–7.

49. Grodsky N, Li Y, Bouzida D, Love R, Jensen J, Nodes B, et al. Structure of the catalytic domain of human protein kinase C beta II complexed with a bisindolylmaleimide inhibitor. Biochemistry. 2006;45(47):13970–81.

50. Leonard TA, Różycki B, Saidi LF, Hummer G, Hurley JH. Crystal structure and allosteric activation of protein kinase C βII. Cell. 2011;144(1):55–66.

51. Stahelin RV, Wang J, Blatner NR, Rafter JD, Murray D, Cho W. The origin of C1A-C2 interdomain interactions in protein kinase Calpha. The Journal of biological chemistry. 2005;280(43):36452–63.

52. Antal CE, Callender JA, Kornev AP, Taylor SS, Newton AC. Intramolecular C2 Domain-Mediated Autoinhibition of Protein Kinase C βII. Cell reports. 2015;12(8):1252–60.

53. Lučić I, Truebestein L, Leonard TA. Novel Features of DAG-Activated PKC Isozymes Reveal a Conserved 3-D Architecture. Journal of molecular biology. 2016;428(1):121–41.

54. Violin JD, Zhang J, Tsien RY, Newton AC. A genetically encoded fluorescent reporter reveals oscillatory phosphorylation by protein kinase C. The Journal of cell biology. 2003;161(5):899–909.

55. Gallegos LL, Kunkel MT, Newton AC. Targeting protein kinase C activity reporter to discrete intracellular regions reveals spatiotemporal differences in agonist-dependent signaling. The Journal of biological chemistry. 2006;281(41):30947–56.

56. Ross BL, Tenner B, Markwardt ML, Zviman A, Shi G, Kerr JP, et al. Single-color, ratiometric biosensors for detecting signaling activities in live cells. eLife. 2018;7.

57. Lu Z, Liu D, Hornia A, Devonish W, Pagano M, Foster DA. Activation of protein kinase C triggers its ubiquitination and degradation. Molecular and cellular biology. 1998;18(2):839–45.

58. Young S, Parker PJ, Ullrich A, Stabel S. Down-regulation of protein kinase C is due to an increased rate of degradation. The Biochemical journal. 1987;244(3):775–9.

59. Hansra G, Bornancin F, Whelan R, Hemmings BA, Parker PJ. 12-O-Tetradecanoylphorbol-13-acetate-induced dephosphorylation of protein kinase Calpha correlates with the presence of a membrane-associated protein phosphatase 2A heterotrimer. The Journal of biological chemistry. 1996;271(51):32785–8.

60. Min X, Zhang X, Sun N, Acharya S, Kim KM. Mdm2-mediated ubiquitination of PKCβII in the nucleus mediates clathrin-mediated endocytic activity. Biochemical pharmacology. 2019;170:113675.

61. Kornev AP, Aoto PC, Taylor SS. Calculation of centralities in protein kinase A. Proceedings of the National Academy of Sciences of the United States of America. 2022;119(47):e2215420119.

62. Di Paola L, De Ruvo M, Paci P, Santoni D, Giuliani A. Protein contact networks: an emerging paradigm in chemistry. Chemical reviews. 2013;113(3):1598–613.

63. Newman M. Networks. Second ed: Oxford University Press; 2018 19 October.

64. Lordén G, Newton AC. Conventional protein kinase C in the brain: repurposing cancer drugs for neurodegenerative treatment? Neuronal signaling. 2021;5(4):Ns20210036.

65. Tovell H, Newton AC. PHLPPing the balance: restoration of protein kinase C in cancer. The Biochemical journal. 2021;478(2):341–55.

66. Takahashi H, Adachi N, Shirafuji T, Danno S, Ueyama T, Vendruscolo M, et al. Identification and characterization of PKCγ, a kinase associated with SCA14, as an amyloidogenic protein. Human molecular genetics. 2015;24(2):525–39.

67. Kazanietz MG, Wang S, Milne GW, Lewin NE, Liu HL, Blumberg PM. Residues in the second cysteine-rich region of protein kinase C delta relevant to phorbol ester binding as revealed by site-directed mutagenesis. The Journal of biological chemistry. 1995;270(37):21852–9.

68. Slater SJ, Ho C, Kelly MB, Larkin JD, Taddeo FJ, Yeager MD, et al. Protein kinase Calpha contains two activator binding sites that bind phorbol esters and diacylglycerols with opposite affinities. The Journal of biological chemistry. 1996;271(9):4627–31.

69. Zheng M, Zhang X, Guo S, Zhang X, Choi HJ, Lee MY, et al. PKCβII inhibits the ubiquitination of β-arrestin2 in an autophosphorylation-dependent manner. FEBS letters. 2015;589(24 Pt B):3929–37.

70. Uphoff CC, Drexler HG. Detection of mycoplasma contaminations. Methods in molecular biology (Clifton, NJ). 2013;946:1–13.

71. Gallegos LL, Newton AC. Genetically encoded fluorescent reporters to visualize protein kinase C activation in live cells. Methods in molecular biology (Clifton, NJ). 2011;756:295–310.

72. Sievers F, Higgins DG. The Clustal Omega Multiple Alignment Package. Methods in molecular biology (Clifton, NJ). 2021;2231:3–16.

73. Waterhouse AM, Procter JB, Martin DM, Clamp M, Barton GJ. Jalview Version 2--a multiple sequence alignment editor and analysis workbench. Bioinformatics (Oxford, England). 2009;25(9):1189–91.

74. Crooks GE, Hon G, Chandonia JM, Brenner SE. WebLogo: a sequence logo generator. Genome research. 2004;14(6):1188–90.

75. Case DA, Betz RM, Cerutti DS, Cheatham III TE, Darden TA, Duke RE, et al. AMBER 2016. University of California, San Francisco; 2016.

76. Scott, Andreas WG, Ross CW. SPFP: Speed without compromise—A mixed precision model for GPU accelerated molecular dynamics simulations. Computer Physics Communications. 2013;184(2):374–80.

77. Csardi G, Nepusz T. The igraph software package for complex network research. InterJournal. 2006;Complex Systems:1695.

